# DIA-NN EasyFilter workflow for the fast and user-friendly critical assessment and visualization of DIA-NN proteomics analysis outcome

**DOI:** 10.64898/2026.03.07.710308

**Authors:** Mabuse Gontse Moagi, Thatiana Ferraz Ferreira, Kristof Endre, Adzani Gaisani Arda, Rini Arianti, Peter Horvatovich, Éva Csösz

## Abstract

Liquid chromatography-tandem mass spectrometry (LC-MS/MS) based proteomics, particularly data-independent acquisition (DIA), has become widely adopted across in One Health approaches for biological and clinical research for quantitative protein characterization. Among the many computational tools available, DIA-NN has demonstrated superior performance; however, the primary output of the current versions is provided as a compact, compressed PARQUET file that can be difficult to interrogate without programming expertise. To address this limitation, we developed DIA-NN EasyFilter (DEF), a fast, user-friendly, KNIME-based workflow for comprehensive protein filtering, and visualization. DEF integrates chromatographic peak-based filtering, curated contaminant libraries, and quantity-quality assessment, along with interactive modules for qualitative and quantitative data exploration. The workflow is optimized for efficient execution within the KNIME local desktop environment and is designed to support end-users in improving accuracy and interpretability without requiring coding skills. We provide detailed description on how to run DEF and demonstrate the utility and robustness of DEF using published large-scale proteomics datasets, showing high comparability across studies regardless of instrument platform or dataset size.

**Table of Contents graphic:** 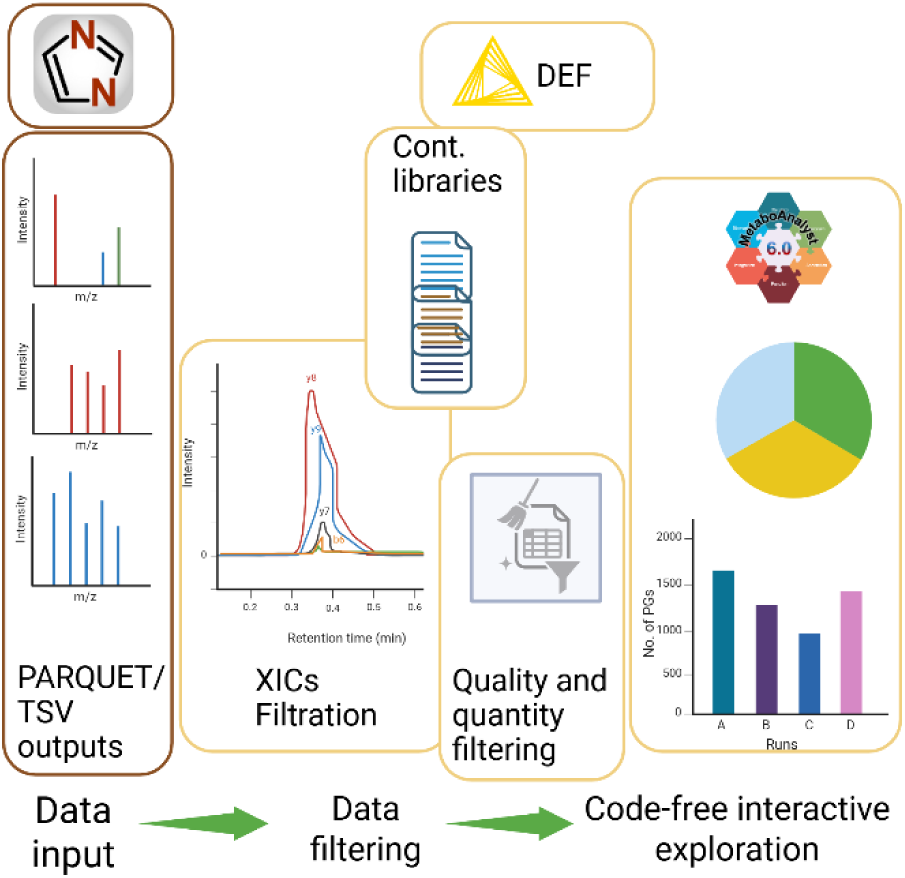

## 1. Introduction

Liquid chromatography-tandem mass spectrometry (LC-MS/MS) based proteomics is widely used to quantify proteins in complex biological and clinical samples^1,2^. In bottom-up workflows, proteins are enzymatically digested and inferred from identified and quantified peptides, which are assembled into protein groups (PGs)^3,4^. To enhance confidence at the protein level, various filtering strategies are commonly employed. For example, some approaches retain only PGs supported by at least two unique peptides^5^, whereas others restrict protein inference to proteotypic peptides derived from predefined peptide lists^6^. Although strategies involving a cut-off of at least two unique peptides improve identification reliability, they may inadvertently reduce identification (ID) depth by excluding PGs that differ from other groups by only a single unique peptide.

Data acquisition is typically performed using data-dependent acquisition (DDA) or data-independent acquisition (DIA)^7^. DDA preferentially fragments the most abundant precursors, introducing sampling bias, whereas DIA fragments all precursors within defined windows^8–10^, improving reproducibility but generating highly multiplexed MS2 spectra that complicate data analysis. Numerous tools have been developed for DDA analysis^11–13^, while DIA requires more advanced peptide-centric or spectrum-centric computational approaches. Examples for spectrum centric tools are DIAUmpire^14^, diaTracer^15^ and DIAmeter^16^, while peptide centric approach has been implemented in Spectronaut^17^, ScaffoldDIA (Proteome Software), EncyclopeDIA^18^, and DIA-NN ^17,19,20^. Among these tools, DIA-NN is widely adopted for accurate identification and quantification using target-decoy scoring and deep neural networks. Version 2.0 introduced the Uncertainty Minimising Solution (QuantUMS) module, which models peptide-level quantification uncertainty from MS1 and MS2 intensities to improve quantitative precision^21,22^.

Despite these advances, systematic filtering and quality assessment of DIA-NN outputs remain challenging, particularly given the PARQUET report format, an industry-standard structure that typically requires dedicated libraries (e.g., PyArrow) for programmatic access. R-based solutions can represent powerful and, in some cases, more streamlined alternatives. Script-based workflows provide full analytical traceability, version control, and high statistical flexibility, particularly when custom models or advanced statistical frameworks are required. However, their effective implementation generally requires proficiency in R, and fully script-based environments may increase the risk of misconfiguration or unintended analytical errors for non-programming users. R-based solutions such as MSDAP^23^ and DIAgiu^24^ primarily focus on quantitative statistics and provide limited support for integrated evaluation of identification performance. In contrast, Skyline^25^ can be used with spectral library to assess the quality of peptide identification and quantification, however its use requires considerable manual analysis and parametrization efforts. Manual inspection is labor-intensive and prone to error, underscoring the need for accessible bioinformatic solutions for users with minimal programming expertise.

Low-code/no-code platforms such as Konstanz Information Miner (KNIME)^26^ and Galaxy^27,28^ facilitate reproducible data analysis through visual workflow designs. Compared with Galaxy, KNIME offers greater flexibility for constructing highly customized, modular workflows through its desktop-based node architecture^29,30^, whereas Galaxy is more optimized for life science analysis.

To alleviate support shortfalls of many open source and free software, KNIME has a community Hub for sharing workflows, components, and extensions. Here users can search for, and access proteomics workflows provided by OpenMS^31^ and the KNIME community. However, no dedicated workflow currently supports comprehensive filtering and evaluation of DIA-NN main reports.

To fill this gap, here, we present DIA-NN_EasyFilter (DEF), a KNIME-based workflow for filtering and analyzing DIA-NN reports (version ≥1.8). DEF enables qualitative and quantitative assessment of peptide and protein identifications without requiring coding skills. The workflow supports two approaches for PG inference, either by 2 unique-peptides or considering proteotypicity, when the proteins are identified based on protein-specific, so called proteotypic peptides^6^. Additionally, when the Extracted Ion Chromatogram (XIC) checkbox is checked in DIA-NN, our workflow offers a peptide quality assessment filter using this report. Large-scale reports (up to tens of millions of rows) can be processed in DEF within minutes, providing an efficient solution in One Health approaches usually dealing with large datasets.

## 2. Methods

### 2.1. KNIME implementation

The DEF workflow was developed (Data S1) using the open-source KNIME Analytics Platform 5.4.3 (www.knime.com). Instructions for downloading and installing DEF (www.knime.com/download/24mabuse/spaces/DEF) are provided in Data S2. All required extensions were installed directly within KNIME, and no external libraries beyond those available in the platform were used.

To enable XIC-based filtration in DEF, the corresponding DIA-NN XIC output must be checked. The DEF workflow is initiated via PARQUET and File Reader nodes, which accept DIA-NN main output (PARQUET or TSV), pg.matrix, and optional XIC reports as inputs. Downstream processing steps are organized into modular sub-workflows (“XICs filter,” “Library choice filter,” “Task selector,” and “Quantity filter”), each configurable through dialog boxes without the need to understand data flows and node functions. In the “XICs filter” sub-workflow, fragment ion features (b- and y-ions) were extracted from the DIA-NN XICs output and concatenated into unique feature sets. Only peptides supported by at least four consecutive b- or y-ions were retained and used to filter the main DIA-NN report. The filtered output was then processed by the “Library choice filter,” which incorporates a curated contaminant list compiled from MaxQuant^32^, cRAP (https://www.thegpm.org/crap/), and the Hao-Group^33^ contaminant datasets. These lists are stored within the workflow data area, however, users may upload a custom contaminant FASTA file as described in Data S2. Matching contaminants and Biognosys iRT peptides were flagged and excluded from the PG identification list.

The “Task selector” sub-workflow was used to split looped tasks, each supporting either a two-unique-peptide rule or a proteotypic peptide-based inference strategy. Under the two-peptide rule, duplicate protein entries were removed and only proteins supported by more than one unique sequence were retained. For the proteotypic strategy, non-proteotypic peptides were filtered analogously, whereas proteotypic peptides required a single unique sequence, and all corresponding unique PGs were retained.

Quality thresholds were defined at the PG level using run-specific PG.Q.Value (≤0.05) and library-level with Lib.PG.Q.Value, and Lib.Q.Value set to ≤0.01. PG.MaxLFQ entries not meeting these criteria were replaced with an arbitrary value of “50,” indicating identified but non-quantified proteins. It should be mentioned that DEF takes quantitative information provided by DIA-NN, using the quantification algorithm implemented in the actual version of DIA-NN (MaxLFQ or QuantUMS in newer DIA-NN versions).

### 2.2. Visualization

Qualitative visualization modules were implemented as composite components to display the number of identified PGs as bar plots. Peptide summary tables were generated to capture both retained and excluded peptides following XIC-based filtration. For quantitative visualization, PG.MaxLFQ values were extracted for each run and used to generate stacked bar plots summarizing run-level quantification performance (Equation 1).

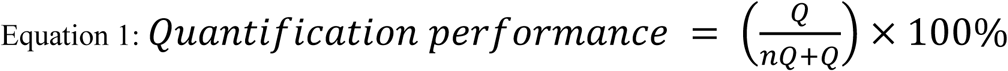

Where:

Q =Number of PGs quantified

nQ = Number of PGs not quantified

Entries not meeting predefined quantity quality thresholds were converted to missing values. These filtered data were subsequently used to construct binary parallel coordinate plots, in which identified PGs were assigned a value of 1 and non-identified PGs a value of 0. A pie chart was further implemented to display the distribution of all PG intensities, including matched contaminants, ordered in descending abundance.

Group-level visualization was implemented within an interactive component supporting comparisons across two to five experimental groups. User-adjustable parameters include missing value thresholds and relative standard deviation (RSD) cut-offs per group. Finally, output tables were reformatted to ensure compatibility with MetaboAnalyst input specifications.

### 2.3. Case studies

To demonstrate the efficiency of our DEF workflow, we performed three separate analyses as Case 1, Case 2 and Case 3, of datasets downloaded from the publicly available ProteomeXchange repository (Table S2) and Case 4 examining results generated in our laboratory.

#### Case 1

Case 1 involved the analysis of PXD029738 dataset that contains raw files obtained from the triplicate analysis of 100 ng loaded peptide originating from Human Embryonic Kidney (HEK) cells^20^. Raw data files were searched against either the author’s provided human FASTA file, DDA-based library, or gas phase fractionation (GPF) generated spectral library by searching the GPF-DIA raw files against the FASTA database in DIA-NN version 2.2 (Table S3). The batch .csv was used to identify the number of proteins from each replicate run. Furthermore, reproducibility between technical replicates was assessed by the median coefficients of variation (CV) using the corresponding PG.MaxLFQ values.

#### Case 2

Case 2 entailed analysis of five replicates from six samples containing 5%, 10%, 13%, 20%, 30% and 40% mouse membrane proteins, respectively, spiked in a yeast background^34^. The authors searched files against a Universal library generated from raw DDA data using a FragPipe, and library-free mode involving in silico library generation. Downloaded DIA-NN (version 1.8.1) TSV output was used as report input in our DEF workflow Batch .csv was used in both modes, and these filtered tables were annotated for organism (mouse or yeast)-specific proteins using Uniprot annotation followed by extraction of mouse-specific proteins. Proteins with two or more missing values, out of five replicates for each sample, were removed and the remaining ones were kept as quantifiable proteins.

#### Case 3

Case 3 included a tri-species mixture sample set (HYE124) composed of human, Escherichia coli, and yeast digests^35^. All AB Sciex dynamic-link library (.dll) files were copied from MSconvert^39^ to the DIA-NN installation directory, which enabled WIFF files to be directly imported into DIA-NN. These WIFF data files were loaded as two batches (TripleTOF 5600 and TripleTOF 6600) and searched against a three-organism library (Table S3). Batch .csv was used and only proteins that were quantified in at least two out of three replicates in one sample were considered. From these, the number of identified proteins and quantification reproducibility expressed in CV, from the effects of mass spectrometers and SWATH acquisition modes, were compared to those reported.

#### Case 4

Case 4 involved the analysis of our data acquired by the examination of three biological replicates from three experimental groups: pre-adipocytes (PA), differentiated white adipocytes (WA), and white adipocytes differentiated in the presence of palmitate (WP) to mimic high-fat conditions. All cells were derived from the subcutaneous adipose tissue specimen of an infant diagnosed with Simpson-Golabi-Behmel syndrome (SGBS)^36^. Differentiation was induced using a standard adipogenic differentiation medium, as described by^37^, for 7 days. For palmitate treatment, WP cells were exposed to 100 µM sodium palmitate in differentiation medium. Following cell harvesting and lysis with 1% SDS, protein concentrations in the lysates were determined using the Pierce™ BCA Protein Assay Kit (Thermo Fisher Scientific). A total of 110 µg of cell lysate per sample was digested using a micro S-Trap™ (ProtiFi, Fairport, NY, USA) column. Proteomic profiling was performed by liquid chromatography-electrospray ionization tandem mass spectrometry (LC-ESI-MS/MS) in DIA mode. The detailed description of the sample preparation and LC-MS analysis can be found in Data S3. Raw data were processed as a batch using DIA-NN 2.2.0 (Table S3).

The resulting PARQUET output files were imported into the DEF workflow, together with the corresponding “XICs” report and “report.pg_matrix” files. Quantification filters were kept at default settings and the HaoGroup contaminant library was applied as this list was most recently updated to include additional contaminants yet excluding unassigned Uniprot entries and human protein standards, unique to MaxQuant and cRAP contaminants. Group-level contrasts were defined within the “Group_Visuals” component in DEF, enabling direct comparison between experimental conditions. In this analysis, three pairs were specified (WA × PA, WP × PA, WP × WA), permitting a maximum of one missing value per replicate, and exported with the “MetaboAnalyst Ready” node.

Subsequent statistical analysis was performed using the MetaboAnalyst 6.0 web-based platform^38^. Within MetaboAnalyst, data were filtered using the low-variance filter, and missing values were imputed using the left-censored method, replacing with one-fifth of the minimum positive intensity value^39^. Data were then median-normalized and log10-transformed. Differential abundance was assessed using a two-sample t-test, with an adjusted p-value < 0.05 and fold-change > 1.5 or fold-change < -1.5 considered to be significant. Principal component analysis (PCA) and volcano plots were generated in MetaboAnalyst. For volcano plots, some non-significant proteins were omitted to simplify the plot.

Protein-protein interaction (PPI) networks, as well as enrichment analyses for biological processes, molecular functions, and KEGG pathways, were constructed for all differentially abundant proteins using the STRING database v12.0^40^. Protein interactions were predicted from all available evidence sources, applying a high-confidence interaction score (0.7) to generate the networks.

## 3. Results and discussion

Our DEF workflow contains several types of components: there are mandatory input components, mandatory yet once-off functions such as “saving location” definer, components for optional and visualization functions. Our workflow connects user defined parameters and splits these into eight branches: (1) and (2) save the filtered output as .csv per sample; (3) and (4) visualize filtered PGs; (5) and (6) visualize filtered protein quantities; and (7) and (8) save the filtered output as .csv of batch samples.

The functionally similar workflow parts were wrapped in components as sub-workflows. These components were further configured to include selection dialogs for simple user interaction. When the researcher chooses to include XICs filtration, all ’b’ or ’y’ ions are concatenated for each peptide, and a small integrated python script search for four consecutive ‘b’ or ‘y’ ion series is defined as inclusion criteria for further analysis^41^ (Table S4). DIA-NN supports XIC visualization through the DIA-NN Viewer and Skyline options. However, although Skyline is freely available, its automated utility directly from DIA-NN is constrained by the requirement to install Skyline and load DIA-NN peptide search results manually.

Mass spectrometry-based proteomics is inherent to contaminants that can be introduced into samples throughout the experimental workflow^42,43^. Common contaminant FASTA files are available for use in various data analysis platforms. The Hao group, however, have postulated that there’s a need for a more frequently updated contaminant list, as some of the contaminant proteins may not be included or are non-contaminant human protein standards^33^. The contaminant sub workflow contains list of three widely used protein contaminant sequence databases, saved into the workflow area, because some contaminants are not universal to all proteomics studies. Contaminants identifications can be used for optimized and tailored protocols and laboratory conditions, and the contaminant list can be obtained from multiple resources such as from MaxQuant, cRAP, and the Hao group. The filter to use a combination of contaminant list or do not use it at all is decided by the user. Users may also upload a custom contaminant FASTA file if their study requires inclusion of non-standard or project-specific contaminant sequences (Data S2).

Recent DIA-NN performs protein quantification with QunatUMS algorithms^22^. By default, our workflow filters columns titled “Quantity” and “PG.MaxLFQ qualities” at 0.5 and 0.7 cutoff, respectively. All PGs that do not meet these criteria are not excluded but are replaced by an imputed ‘nan’ value for its quantity. The further filtering of quantified PGs is left at user decision.

### Case studies

To verify the use of our DEF workflow, we downloaded .raw files (Case 1), DIA-NN TSV output (Case 2), and WIFF files (Case 3), from a public repository. Platform and DIA-NN parameter details are given in Table S2 and Table S3, respectively^20,35^.

Re-analysis of the data in Case 1 with DEF workflow provided over 4200 PGs which is similar to those reported in the original publication (Table 1).

**Table 1.** Comparison of the number of protein identifications in single injections from the DDA-, FASTA-, and GPF-based strategies.

|  | GPF |  | FASTA |  | DDA |  |
| --- | --- | --- | --- | --- | --- | --- |
|  | Jiang et al.<sup>20</sup> | DEF | Jiang et al.<sup>20</sup> | DEF | Jiang et al.<sup>20</sup> | DEF |
| HEK293_100ng_replicate_1 | 4752 | 4796 | 4691 | 4647 | 4270 | 4433 |
| HEK293_100ng_replicate_2 | 4809 | 4863 | 4666 | 4689 | 4253 | 4506 |
| HEK293_100ng_replicate_3 | 4959 | 4936 | 4766 | 4722 | 4467 | 4612 |

Of the three replicates, PGs with no missing values were retained to evaluate the number of quantified proteins and quantitative reproducibility based on median and distribution of CV between replicates. As expected, the numbers of reported proteins are likely to be different due to a different filtering approach. However, a similar trend could be observed from our KNIME workflow output to that of the original publication (Figure 2).

**Figure 1.**
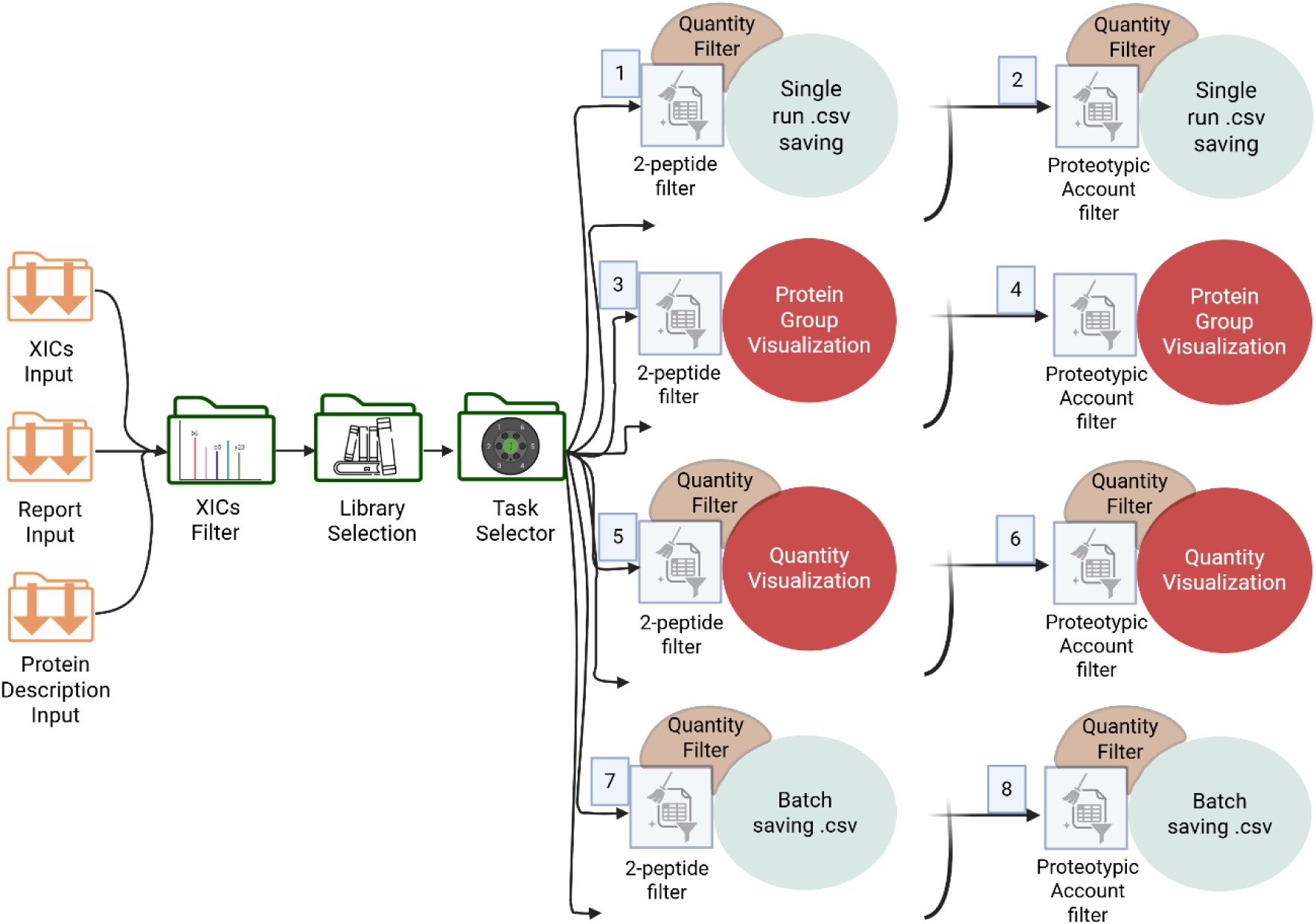
Workflow illustration showing input (orange), configuration components (green), optional filters (brown), and visual (red) components. A more detailed version of the figure can be found as Figure S1.

**Figure 2.**
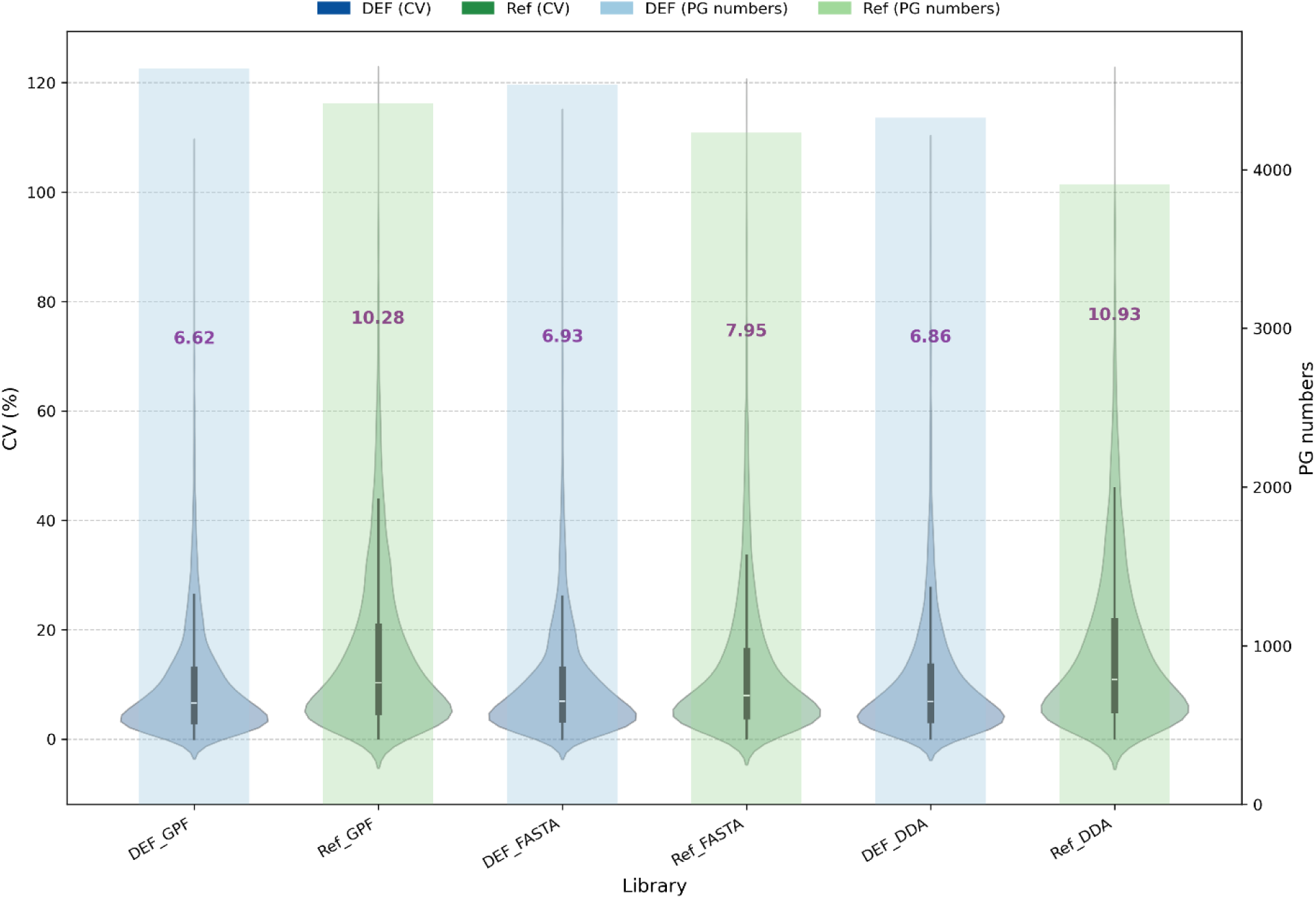
Quantification accuracies from a GPF-, FASTA-, and DDA-based library search. DEF highlights results filtered in our workflow with an additional XIC filtration requiring consecutive b and y ions, while Ref represents those from the original publication. Bars (right y-axis) indicate the number of quantified PGs per library, while the overlaid violins (left y-axis) illustrate the distribution of their coefficients of variation (CV, %). Median CV values are annotated above each violin.

DDA-based method resulted in slightly fewer number of proteins identified compared to FASTA- and GPF-based libraries. Additionally, these different strategies showed comparable quantitative reproducibility, with median CV values of 6.62%, 6.93%, and 6.86% corresponding to GPF, FASTA, and DDA-based libraries, respectively. The slight improvements in the number of protein and median CV values when comparing our DEF workflow to the Case 1 publication could be attributed to the QuantUMS algorithms implemented in recent DIA-NN version 2.2^44^ to earlier DIA-NN version used by the authors.

For Case 2 we tested the use of TSV output from DIA-NN version 1.8.1. The downloaded reports were composed of five replicates analyzed on Thermo QE HF and Bruker timsTOF Pro with a universal and an in silico library. The number of identified PGs by DEF based on the proeotypic rule is similar to the razor protein inference approach^45^ which was used in the original publication. A comparable number of mouse protein identifications was observed on data from both types of instruments and using different library generation strategies (Table 2).

**Table 2.** The number of identified mouse PGs from both Thermo QExactive HF (HF) and timsTOF Pro (TIMS) instruments when searched against a universal library and DDA-independent library (Library free)

|  | Universal library |  | Library free |  |
| --- | --- | --- | --- | --- |
|  | Lou et al.<sup>34</sup> | DEF | Lou et al.<sup>34</sup> | DEF |
| HF | 4923 | 4903 | 5186 | 5166 |
| TIMS | 7128 | 7176 | 6616 | 6664 |

To compare quantification performance, we exported the result file in .csv format from the batch samples branch. Although enabling quantify filters is recommended, as no filters were used by authors, we opted not to use any quantity filters in DEF to achieve comparable results with the one reported in the original publication. Mouse PGs with at least two missing values out of the five replicates per condition were retained as quantifiable PGs. The different library types and MS platforms analysed by our workflow showed high similarities to those reported in the original publication (Figure 3). However, our workflow filters showed slight improvements in reproducibility of mouse PG quantities, with median CVs of 4.35-10.28% as opposed to 4.43-11.84% reported in^49^.

**Figure 3.**
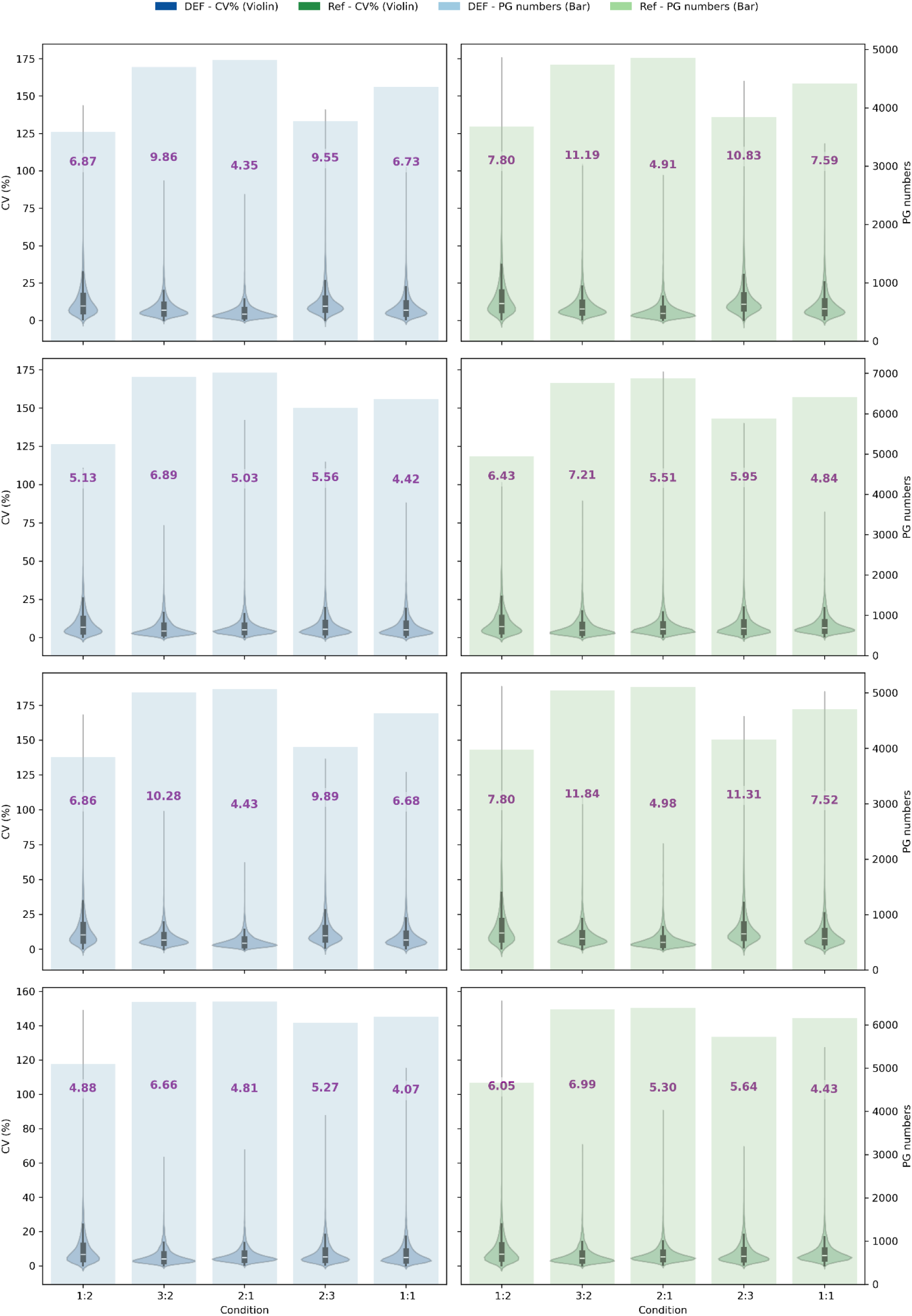
Comparative evaluation of quantification performance showing DIA-NN_ EasyFilter (blue) and reported in reference publication (green) for Oritrap HFX and timsTOF Pro instruments and various library generation methods (universal and library free). Bar plots (right axis) represent the number of quantified PGs, while overlaid violin and box plots (left axis) show the distribution of their coefficients of variation (CV, %). The median CV value for each group is indicated above the violins.

Case 3 was aimed at evaluating the quantification and identification performance in function of the mass spectrometer model and the number of used SWATH acquisition windows (32 fixed and 64 variable windows) compared to those reported in the original publication. The authors followed a two-steps iterative process. In the first round, the developers analyzed the datasets using the most recent public versions of their tools with optimized settings. This was followed by an open discussion with the developers, who subsequently refined their software to address all identified limitations. These improvements were then assessed in a second validation round (Table 3). A slightly higher number of proteins were identified from DEF outputs as compared to those reported. This was expected as several studies have shown that, in terms of identification rate, and quantification performance, DIA-NN outperforms Spectronaut, OpenSWATH, Skyline^46^, and DIA-Umpire^47^.

**Table 3.** Numbers of reported proteins, quantified in at least 2 replicates out of 3, identified by different software tools and DEF in HYE124 cell lines analysed by two types of Sciex QTOF instruments and number and types of SWATH windows^39^.

|  |  | Number of proteins in HYE124 (iteration 2)<sup>35</sup> |  |  |
| --- | --- | --- | --- | --- |
|  |  | 32 fixed | 64 variable | Difference |
| OpenSWATH<sup>35</sup> | 5600 | 3,339 | 4,086 | +22% |
|  | 6600 | 4,035 | 4,636 | +15% |
|  | Difference | +21% | +13% |  |
| SWATH 2.0<sup>35</sup> | 5600 | 2,125 | 3,119 | +47% |
|  | 6600 | 2,656 | 3,946 | +49% |
|  | Difference | +25% | +27% |  |
| Skyline<sup>35</sup> | 5600 | 2,878 | 3,629 | +26% |
|  | 6600 | 4,046 | 4,692 | +16% |
|  | Difference | 41% | 29% |  |
| Spectronaut<sup>35</sup> | 5600 | 3,332 | 3,669 | +10% |
|  | 6600 | 4,226 | 4,675 | +11% |
|  | Difference | +27% | +27% |  |
| DIA-Umpire<sup>35</sup> | 5600 | 1,566 | 1,781 | +14% |
|  | 6600 | 2,729 | 3,673 | +35% |
|  | Difference | +74% | +106% |  |
| DEF | 5600 | 4873 | 5386 | +11% |
|  | 6600 | 5658 | 6707 | +19% |
|  | Difference | +16% | +25% |  |

Our workflow data shows how the 64 variable (64var) SWATH window setup resulted in more proteins when compared to the 32 fixed (32fix) window setup. However, the instrument type exerted a larger impact on protein identifications as shown on Table 3. These results were in agreement with those reported in the original publications, where SWATH window setup changes from 32fix to 64var resulted in a 9-37% increase, yet 14-102% more protein identifications were observed for data acquired with TripleTOF 6600 compared to TripleTOF 5600 system^35^. Similarly to Cases 1 and 2, a high technical reproducibility was observed among replicates, and these were in accordance with those reported in the original articles (Figure S2, Table 4). These results show that our DEF workflow ensures rapid and accurate post processing of DIA-NN output, completing a 35-sample batch in under 14 minutes (Table S5).

**Table 4.** Quantification reproducibility evaluated by means of a median coefficient of variation (CV%).

|  | TripleTOF<br>5600 32fix | TripleTOF<br>5600 64var | TripleTOF<br>6600 32fix | TripleTOF<br>6600 64var |
| --- | --- | --- | --- | --- |
| OpenSWATH<sup>35</sup> | 6% | 5% | 8% | 6% |
| SWATH2.0<sup>35</sup> | 5% | 4% | 8% | 6% |
| Skyline<sup>35</sup> | 6% | 6% | 9% | 5% |
| Spectronaut<sup>35</sup> | 4% | 4% | 3% | 3% |
| DIA-Umpire<sup>35</sup> | 10% | 10% | 12% | 12% |
| DEF | 7% | 9% | 6% | 7% |

In Case 4 study we used our workflow to identify and quantify proteins that change during adipocyte differentiation from preadipocytes (PA) to mature white adipocytes in the presence (WP) or absence of palmitate (WA). The number of identified proteins is presented in Table 5, while the lists of identified and quantified proteins are available in Table S6. As expected, applying a 2-peptide inference requirement reduced the number of identified PGs. Even so, more than 300 PGs were confidently identified per replicate under the 2-peptide rule, and over 500 PGs when restricting identification to proteotypic peptides.

**Table 5.** Number of identified PGs in Case 4 study. Number of PGs with a 2-peptide and proteotypic inference rule searched against a human FASTA and combined HaoGroup contaminant list are shown in case of each sample. PA is for pre-adipocytes, WA stands for differentiated white adipocytes, WP for white adipocytes differentiated in the presence of palmitate and Rep for replicates.

|  | Number of PGs |  |
| --- | --- | --- |
|  | 2-peptide rule | Proteotypic rule |
| PA_rep1 | 566 | 754 |
| PA_rep 2 | 399 | 535 |
| PA_rep 3 | 416 | 554 |
| WA_rep 1 | 588 | 787 |
| WA_rep 2 | 706 | 907 |
| WA_rep 3 | 628 | 830 |
| WP_rep 1 | 627 | 813 |
| WP_rep 2 | 696 | 904 |
| WP_rep 3 | 684 | 871 |

Additionally, we can see the effect of XICs filtration on identified PGs number (Figure S3). The ‘Included peptides table’ also contains a ‘Modifications’ column per run that extracts the type of modification and location on each peptide by showing the modified amino acid preceding the modification annotation (Table S7). The ‘Quantity visuals’ composite provides an interactive pie chart for a specific threshold for sample plot. This threshold defines a cut-off for chart sectors with protein description, values below cut-off will remain in the ‘other’ segment of the pie chart. This chart displays all proteins, including contaminants, before filtration within the loop body. Although these are filtered out and are not included in exported results from DEF, we aimed to enable users to visualize which PGs had high intensities (Figure S4). Chromatogram intensity plays a major role in mass spectrometry-based proteomics studies, this information often allows users to judge if a sample contains specific peptides^45^. The Pie chart frequencies represent scaled DIA-NN PG.MaxLFQ values, which inputs aggregated peptide intensities across all samples. This in turn enables the user to see how much contaminants contributed to the overall intensity per sample and if these may have suppressed the detection of low abundant peptides^33^.

To evaluate quantification performance from complementary perspectives, three plots can be generated. The first is a parallel coordinate plot using a binary encoding in which identified PGs were assigned a value of 1 and missing PGs in a particular sample with a value of 0, emphasizing patterns of PGs identification across samples and enabling rapid assessment of PGs detection consistency (Figure 4A and 4B). The second parallel coordinate plot incorporated min-max-scaled (0-1) PG quantities, highlighting relative abundance patterns independent of absolute signal intensity and providing a more detailed view of PG-specific trends across samples and experimental conditions (Figure 4C and 4D). In addition, a bar graph summarizing the percentage of quantified PGs for each run offers a straightforward comparison of PGs quantification completeness across datasets (Figure 4E and 4F). Furthermore, a summarized PG quantity table displaying the total abundance of each PG across all runs can be visualized and exported (Table S6). Selecting rows within this table dynamically highlights the corresponding PG in both parallel coordinates plots, enabling a multi-layered assessment of PGs detection coverage and quantitative behavior across samples (Figure 4A and 4B).

**Figure 4.**
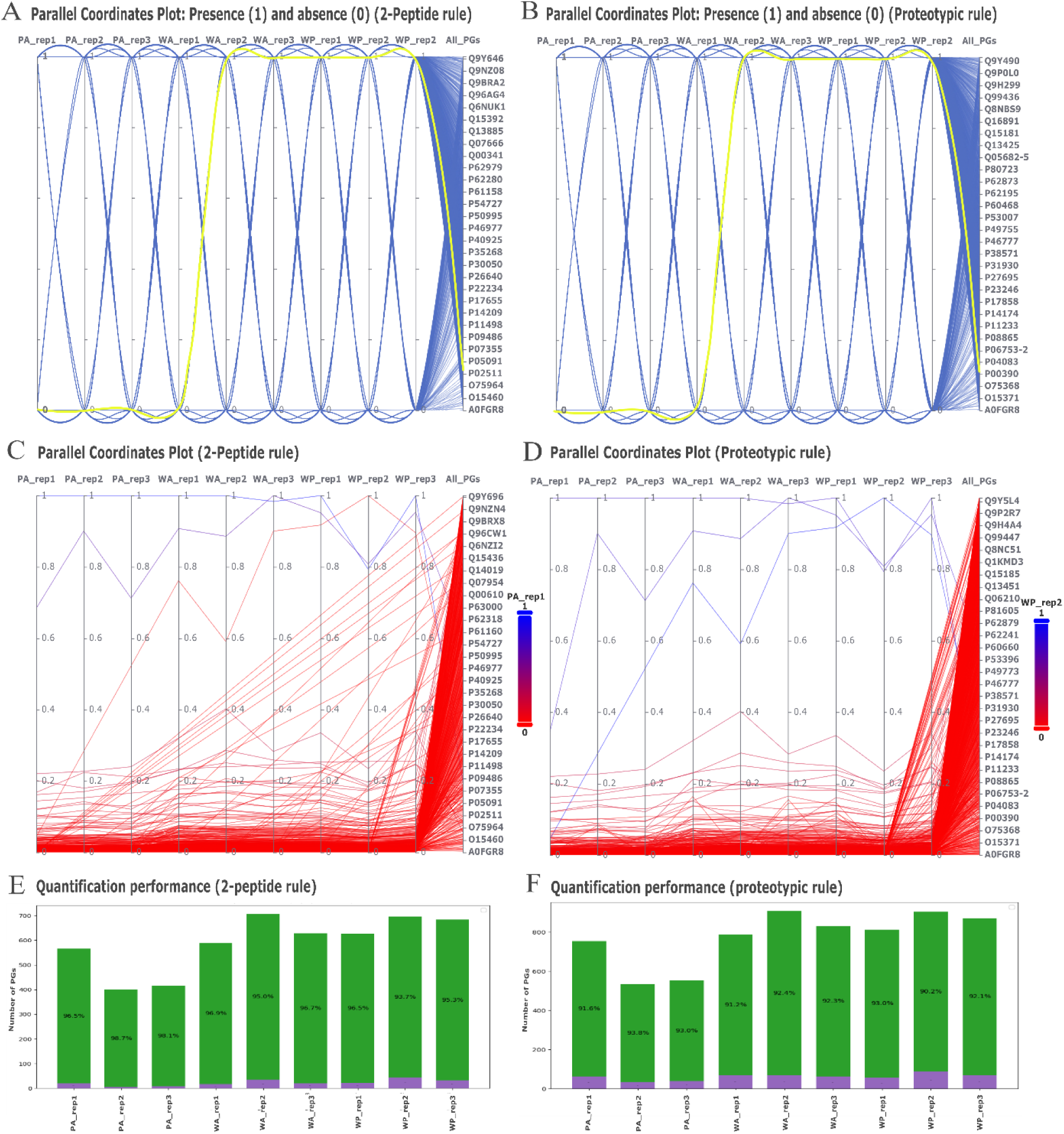
DEF workflow parallel coordinate plots showing a run-level comparison of protein identifications using 2-peptide rule (A) and proteotypic rule (B) with the arbitrarily selected O60240 protein (yellow line). DEF workflow parallel coordinate plot showing a run-level comparison of scaled protein quantities for a 2-peptide rule (C) and proteotypic rule (D) where the intensity dimension was color-coded according to user defined runs (in this case WP_rep2 for panel C and PA_rep1 for panel D). In panels A and B, the left y-axis indicates binary identification status (absence = 0; presence = 1), whereas in panels C and D it represents scaled PG intensities. The right y-axis in panels A-D displays a subset of PG identifiers. Panels E and F show the ratio of sample specific quantifiable PGs obtained using the two-peptide rule (E) and the proteotypic rule (F). The x-axis represents individual runs, and the y-axis indicates the number of PGs expressed as a percentage, with quantified PGs shown in green and non-quantified PGs in purple.

For examining the differentially abundant proteins (DAPs) between the groups, Metabo-Analyst was applied. Within MetaboAnalyst, PGs that remained nearly constant across compared conditions were excluded based on an interquartile range (IQR) threshold of 10% or 25%, depending on whether the total number of PGs was below or above 500, respectively. This was followed by normalization (Figure S5) where log10 PGs intensities within a sample were scaled to the same median value. To further characterize the expression patterns across the sample types, PCA was performed (Figure S6). As expected, the PA (pre-adipocytes) group displays a more distinct pattern compared to the differentiated white adipocytes (WA and WP). In contrast, WA and WP (Figure S6E and S6F) exhibited highly similar patterns, suggesting that the adipocyte differentiation process was the most significant factor driving the observed changes in the proteomic profiles. To better understand the proteomic alterations associated with adipocyte differentiation and palmitate treatment, volcano plots were generated to illustrate the distribution and magnitude of differentially expressed proteins across the sample groups (Figure S7). From DEF workflows obtained with the two types of protein inference, statistical summaries were compiled in MetaboAnalyst (Table 6). Here, p-values are calculated using two-sample t-tests, and differential abundance was determined using a log2 fold-change (log2FC) threshold of 1.5 with significance set at Benjamini-Hochberg FDR q-value of < 0.05 (Table S8). The analysis of protein abundance across different adipocyte conditions revealed 43 more abundant and 72 less abundant proteins in the WA compared to PA groups using the 2-peptide rule. Using proteotypic rule PGs inference, 57 proteins showed higher and 69 lower abundance in WA group. For the WP vs. PA comparison, 49 proteins were more- and 83 less abundant in WP compared to PA using the 2-peptide rule, and 66 PGs showed higher and 95 lower abundance were observed using the proteotypic rule protein inference. Between the WA and WP groups 3 proteins showed statistically significant change. Under the 2-peptide rule cytochrome c oxidase subunit 2 (MT-CO2, P00403) was found to be less abundant, while serotransferrin (TF, P02787) and 60S ribosomal protein L10 (RPL10, P27635) were more abundant in WP. Cytochrome c oxidase subunit 2 is linked to oxidative phosphorylation being part of the mitochondrial electron transport chain. Serotransferrin, in turn, is responsible for iron transport, while the 60S ribosomal protein L10 acts as a component of actively translating ribosomes^40^. According to the proteotypic rule, only the serotransferrin was more and the cytochrome c oxidase subunit 2 less abundant for WP compared to WA.

**Table 6.** A summary of sample group-level DEF output and statistics as performed in MetaboAnalyst showing the number of identified proteins, differentially abundant proteins (DAP) and significantly increased and decreased proteins.

| DEF |  |  | Statistics (MetaboAnalyst) |  |  |
| --- | --- | --- | --- | --- | --- |
| Inference rule | Group | Number of proteins | DAP | FC↑ | FC↓ |
| 2-peptide | WA x PA | 448 | 115 | 43 | 72 |
|  | WP x PA | 448 | 132 | 49 | 83 |
|  | WP x WA | 640 | 3 | 2 | 1 |
| Proteotypic | WA x PA | 601 | 126 | 57 | 69 |
|  | WP x PA | 601 | 161 | 66 | 95 |
|  | WP x WA | 860 | 2 | 1 | 1 |

To gain functional insights into the differentially expressed proteins, Biological Process, Molecular Function, as well as KEGG pathways enrichment analyses were performed (Figure S8). Between the WA×PA and WP×PA comparisons of PGs obtained with the 2-peptide rule, the biological processes for actin cytoskeleton organization, regulation of body fluid levels, supramolecular fiber organization, negative regulation of molecular function, regulation of apoptotic process, and generation of precursor metabolites and energy were commonly enriched in both WP×PA and WA×PA comparisons. Consistent with these, molecular function enrichment analysis revealed a predominance of remodeling activities, such as actin binding, actin filament binding, and structural constituents of the cytoskeleton for both WP×PA and WA×PA comparisons. This shared signature indicates that the transition from precursor cells to the white adipocyte state involves conserved structural remodeling of the cytoskeleton and supramolecular assemblies, accompanied by enhanced oxidoreductase activity. Since just a few proteins were found to be differentially abundant in the palmitate treatment, pathway enrichment analyses for the WP×WA group were not possible.

Finally, in the comparison between mature adipocytes treated or untreated with palmitate and preadipocytes, both inference rules identified KEGG’s enriched pathways primarily associated with carbon metabolism and metabolic pathways. It suggests a redirection of carbon flux toward lipogenic processes and the activation of multiple anabolic routes that can support triglyceride synthesis and energy storage. Notably, proteoglycans in cancer were specifically enriched in palmitate treated adipocytes under the proteotypic rule, which may indicate that palmitate exposure can trigger remodeling of extracellular matrix components in response to fatty acid-induced stress.

## Limitations and future perspectives

While DEF provides a structured and user-friendly framework for DIA-NN post-processing within the KNIME environment, several limitations should be acknowledged. First, although the workflow is suitable for medium-scale studies, as in Case 2 with a 35-sample run, its scalability may be limited compared to web-based such as Galaxy, which are designed to support large cohort analyses. DEF local execution may therefore require memory optimization when applied to very large datasets.

Second, DEF is currently tailored specifically to DIA-NN output formats. Although this ensures seamless integration, it limits direct applicability to datasets processed with alternative DIA search engines.

Finally, while DEF emphasizes standardized filtering, protein inference logic, and quality control, it does not currently implement advanced or alternative quantification algorithms beyond those provided by DIA-NN. Future development will therefore focus on expanding compatibility to a broader range of input formats and raw data sources, as well as incorporating additional quantification and normalization strategies to enhance flexibility and analytical depth.

## Conclusion

In conclusion, our DEF workflow combines data exploration, retrieval, and analysis with the aid of flow controls. The advantage of DEF over other data analysis solutions is that the interactive visual workflow provides a simple readjustment of the analysis parameters and visualization of the results with interactive graphs. Across three case studies, DEF produced identification and filtering outcomes comparable to those reported in the original studies, demonstrating that the workflow can reliably reproduce established filtering strategies while simplifying the post-processing of DIA-NN outputs in a short period of time. This allows researchers with less solid background in programming to critically assess DIA-NN output in PARQUET format in detail, filling a critical gap in the end user community of this community-wide used LC-MS/MS DIA analysis. The XICs of four consecutive b and y ion series visualization and inclusion/exclusion of identified peptides, the availability of 3 different contaminants lists with option to include user specific contaminants, and having the option to filter protein quantitative data with protein inference obtained by the 2-unique peptides or proteotypicity rules enables researchers to tailor their workflow that will best fit for their particular studies. Although Galaxy integrates better with existing bioinformatics workflows, KNIME is better for rapid workflow development and can be implemented without extensive programming background.

As discussed above, DEF is currently optimized for DIA-NN output (v.1.8 and above) formats and operates within a local KNIME environment. Nevertheless, for small- to medium-scale studies, often produced in clinical and One Health studies, the workflow enables efficient and reproducible evaluation of complex LC-MS/MS DIA datasets without requiring coding or KNIME Analytics platform expertise. We are confident that our step-by-step utility guide (Data S2), will enable researchers with no programming background to efficiently use sophisticated data analysis approaches for the analysis and assessment of their proteomics data.

## Data and tool availability

Our workflow can be downloaded from GitHub (https://github.com/MabuseM/DIA-NN_EasyFilters_with_KNIME) or KNIME Hub (https://hub.knime.com/24mabuse/spaces) and executed locally. A detailed protocol on how to set up the workflow is presented in supplementary Data S2. The mass spectrometry proteomics data have been deposited to the ProteomeXchange Consortium via the PRIDE^48^ partner repository with the dataset identifier PXD072043.

## Supporting Information

- Technical Implementation of the DEF Workflow (Data S1); Step-by-step KNIME-based DEF DIA-NN_EasyFilters_(DEF) workflow set up and usage (Data S2); Sample preparation (Data S3); Compacted high resolution DEF workflow (Figure S1); Case 3 DEF filtered quantification reproducibility (Figure S2); DEF qualitative evaluation (Figure S3); Case 4 DEF protein group intensities chart (Figure S4); Case 4 MetaboAnalyst median normalization visualization (Figure S5); Case 4 WA-PA, WP-PA, and WP-WA MetaboAnalyst PCA plots (Figure S6); Case 4 WA-PA, WP-PA, and WP-WA MetaboAnalyst volcano plots (Figure S7); Case 4 WA-PA, and WP-PA biological process, functional and KEGG pathway enrichment (Figure S8) (PDF)
- MaxQuant, cRAP, and Hao Group contaminant lists (Table S1) (XLSX)
- Case 1 - 3 ProteomeXchange file details (Table S2); Case 1,3, and 4 detailed DIA-NN 2.2 parameters (Table S3); DEF workflow run-time evaluation (Table S5); Case 4 peptide summary (Table S7); Case 4 Differentially abundant proteins (DAP) (Table S8) (XLSX)
- Case 1, 3 and 4 b- and y-ion features (Table S4) (XLSX)
- Case 1 - 4 DEF group-level outputs (Table S6)

## AUTHOR INFORMATION

### Author Contributions

Conceptualization, M.G.M. and É.C.; methodology, M.G.M.; software, M.G.M.; formal analysis, M.G.M., T.F.F, K.E., A.G.A., and R.A.; investigation, K.E., A.G.A., and R.A.; resources, É.C.; data curation, M.G.M., T.F.F, K.E., A.G.A., and R.A.; writing—original draft preparation, M.G.M., P.H and T.F.F.; writing—review and editing, É.C., and P.H.; visualization, M.G.M. and T.F.F; supervision, É.C.; project administration, É.C.; funding acquisition, É.C., K.E., A.G.A., and R.A. All authors have read and agreed to the published version of the manuscript.

### Funding Sources

This work was funded by NKFIH - FK145866, NKFIH-PD146202, DHET Academic Support Funding (ASF), János Bolyai Research Scholarship of the Hungarian Academy of Sciences funds and Tempus Public Foundation.

## ACKNOWLEDGMENT

K.E acknowledges Prof. Dr. Pamela Fischer-Posovszky and Prof. Dr. Martin Wabitsch (Department of Pediatrics and Adolescent Medicine, University Medical Center Ulm, Ulm, Germany) for providing SGBS adipocytes. We thank the help of Dr. Gyula Hoffka for the critical review of the manuscript.

## ABBREVIATIONS

LC-MS/MS: liquid chromatography-tandem mass spectrometry
DIA: data-independent acquisition
DEF: DIA-NN_EasyFilter
KNIME: Konstanz Information Miner
PG: protein group
DDA: data dependent acquisition
NNs: neural network
PTM: post-translational modifications
QuantUMS: quantification method using an Uncertainty Minimising Solution
LCAPs: low-code/no-code data science and integration platforms
APIs: application programming interfaces
XICs: extracted ion chromatogram
RSD: relative standard deviation
HEK: Human Embryonic Kidney
GPF: gas phase fractionation
CV: coefficients of variation
dll: dynamic-link library
PA: pre-adipocytes
WA: differentiated white adipocytes
WP: white adipocytes differentiated in the presence of palmitate
SGBS: Simpson-Golabi-Behmel syndrome
PCA: principal component analysis
PPI: protein-protein interaction
IQR: interquartile range
PCA: principal component analysis
DAP: differentially abundant proteins

## Notes

### Competing Interest Statement

The authors have declared no competing interest.

