## Supplementary material for "DIA-NN EasyFilter workflow for the fast and user-friendly critical assessment and visualization of DIA-NN proteomics analysis outcome": Data s2

### Supplementary Data

#### Data S2: Step-by step tutorial on how to set up the KNIME-based DEF workflow

##### S2.1. Getting Started (Figure DS2-1)

Download and install the KNIME Analytics Platform (www.knime.com/download) and the DEF workflow from the KNIME Hub (https://hub.knime.com/24mabuse) or https://github.com/MabuseM/DIA-NN_EasyFilters. After installation, open KNIME, switch to the Classic user interface, and import the DEF workflow.

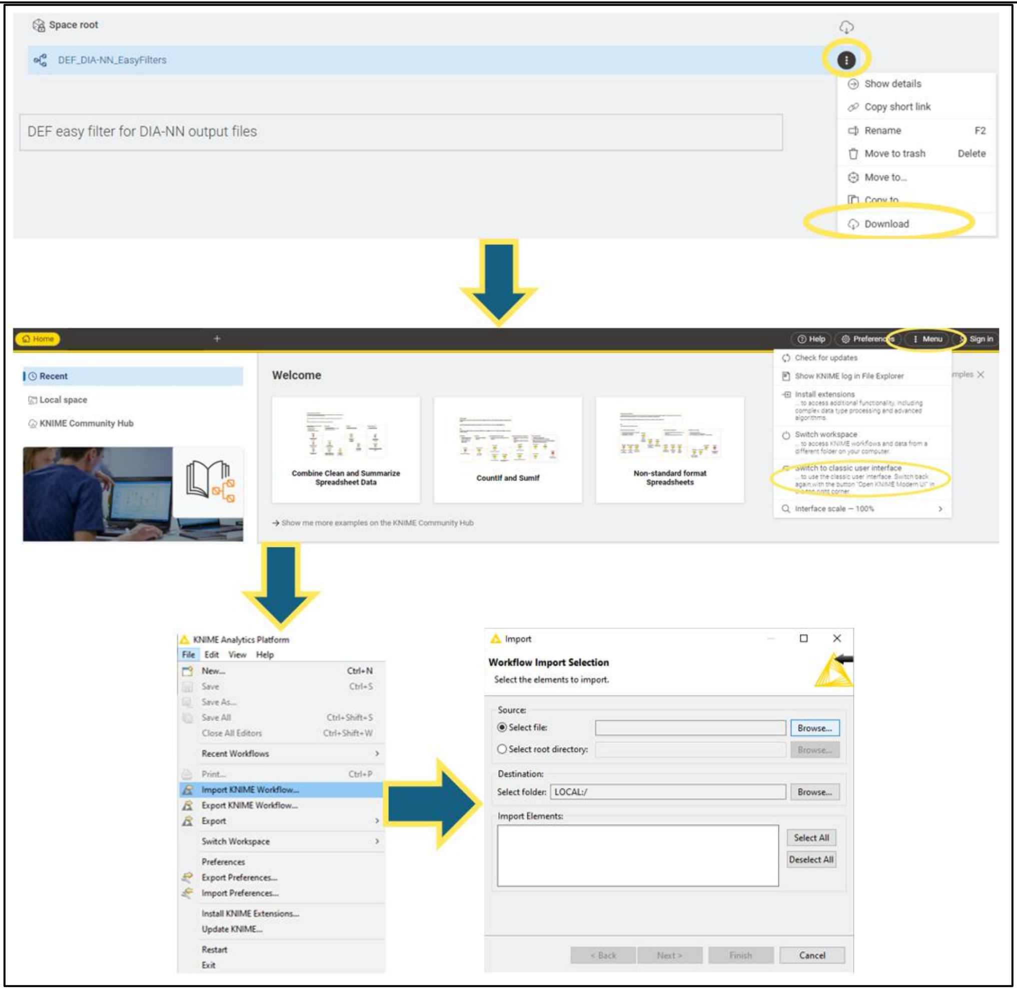

Figure DS2-1: Getting Started

##### S2.2. Extension installation (Figure DS2-2)

Open the DEF workflow and install all required extension when prompted. After restarting, the workflow is ready for use. You may switch back to the Modern UI for a simplified interface.

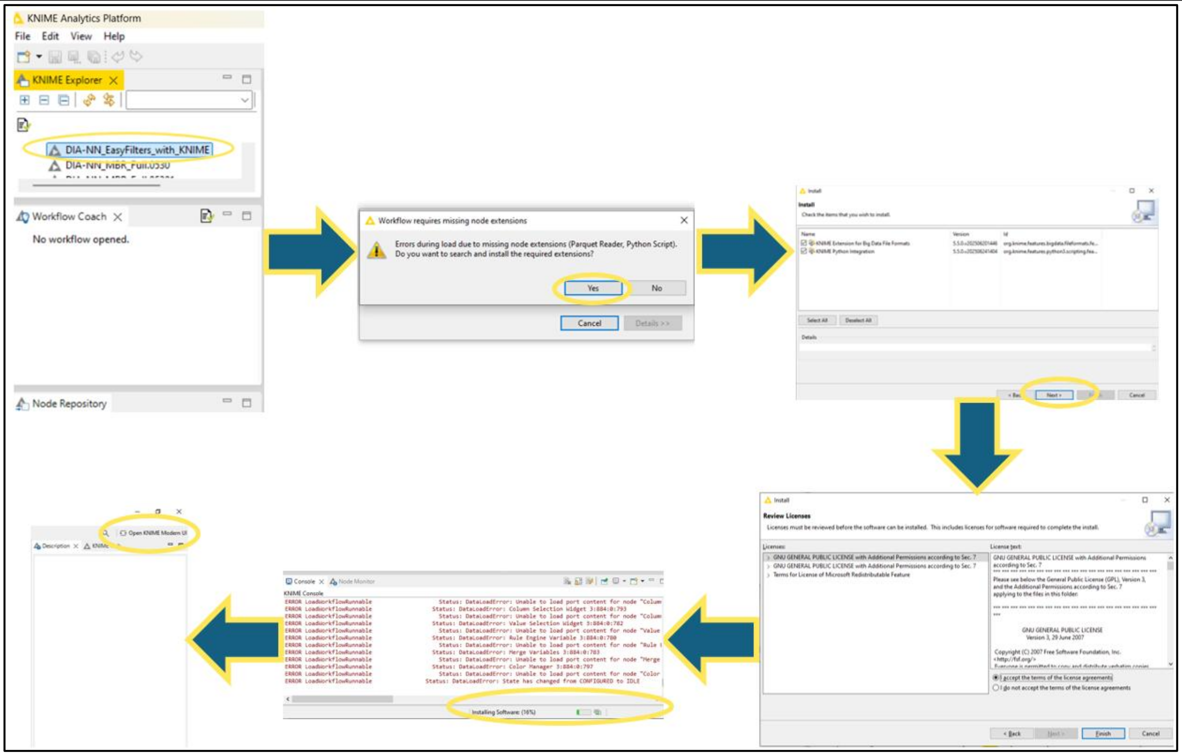

Figure DS2-2: Extension installation

S2.3. <u>Loading DIA-NN Reports (Figure DS2-3)</u>

- Open the *PARQUET Reader* node and select the DIA-NN main report (PARQUET format) from your local file system.
- If XIC filtering is used, open the corresponding PARQUET Reader node and select the report_xic folder.
- Open the *File Reader* node and select the “report.pg_matrix.tsv” file. Click **Auto Detect** to ensure correct TSV formatting.

For compatibility with different DIA-NN versions:

- When using a TSV main report, load “Parquet_reader_Blank” in the PARQUET Reader node.
- When using a PARQUET main report, load “TSV_reader_Blank” in the File Reader node.

Define the output directory in the *Create File/Folder Variables* node by selecting a base location for all DEF outputs. This step is required only once unless the output path is changed.

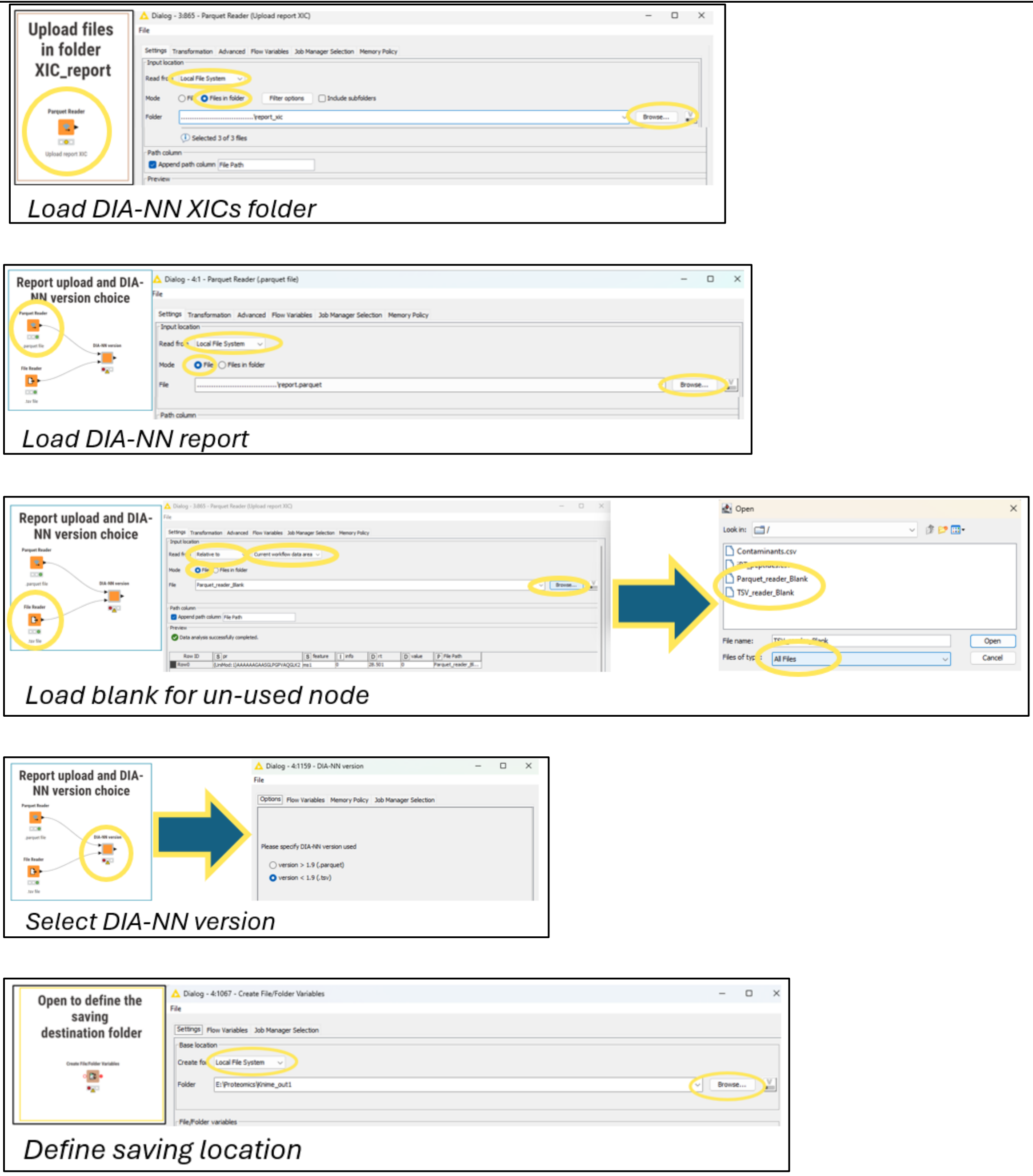

Figure DS2-3: Loading DIA-NN Reports

#### S2.4. <u>Configuring Components (Figure DS2-4)</u>

Adjust workflow parameters within the component dialog boxes, including:

- XIC filtering (on/off)

- Contaminant library selection
- Processing branch (task selection)
- Quantity quality thresholds

*Troubleshooting:* If configuration options are not visible, execute and immediately cancel the component node to refresh the dialog interface.

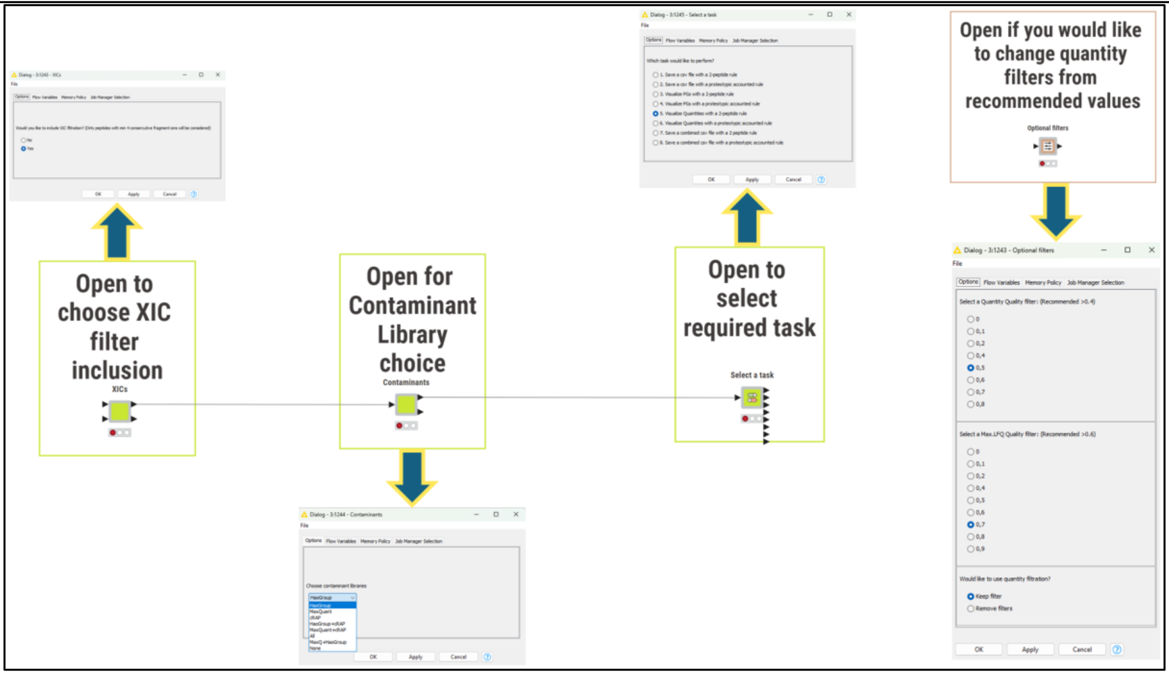

Figure DS2-4: Configuring Components

S2.5. <u>Uploading a Custom Contaminant FASTA File (Optional) (Figure DS2-5)</u>

If a custom contaminant FASTA file is to be used, open the "Contaminants" component within the workflow. In the "*Case Switch Start*" node, change the active port from 1 (default internal list) to 0 (custom input).

Next, open the *FASTA Reader* node and upload the desired FASTA file from your local file system. Use the *Cell Splitter* nodes to extract the protein identifier from the FASTA header based on the appropriate delimiters (e.g., "|" and "_", depending on the header format). Retain only the column containing the protein identifier, remove all other columns, and rename the remaining column to "ID" to ensure compatibility with downstream filtering steps.

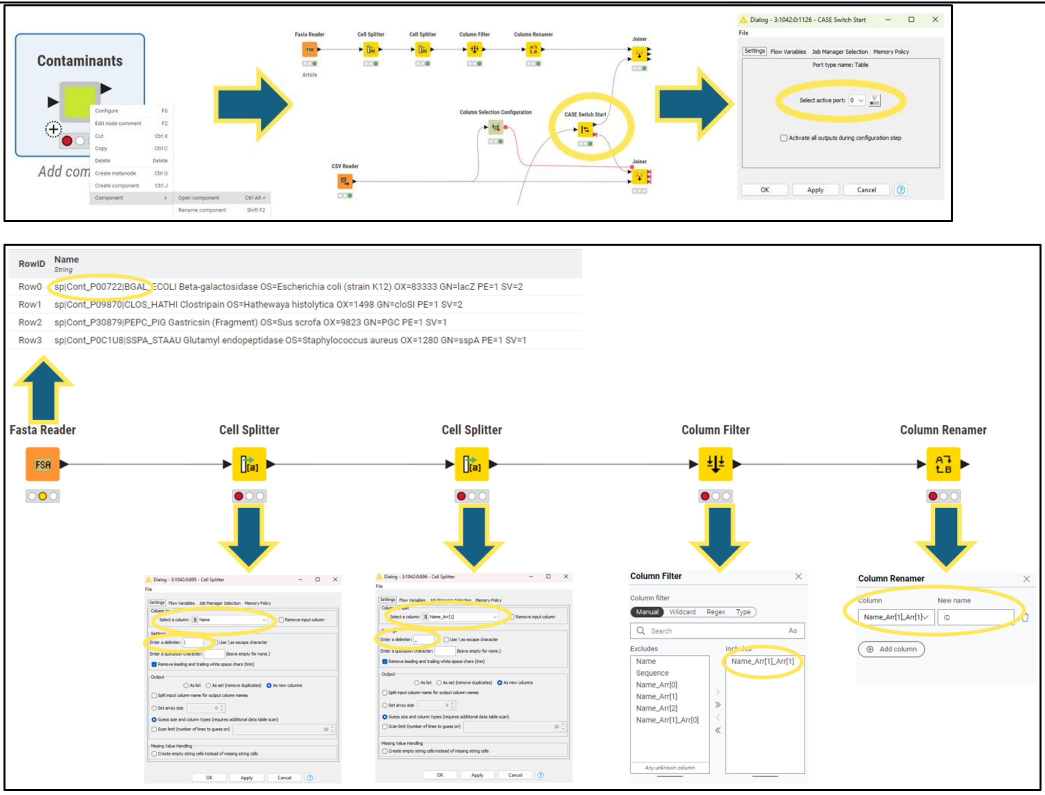

Figure DS2-5: Uploading a Custom Contaminant FASTA File

#### S2.6. <u>Workflow Execution (Figure DS2-6)</u>

Execute the workflow using "Execute All" (Shift + F7) or run individual nodes (F7). Hovering over a node reveals the execution icon, allowing node-level execution including all upstream steps.

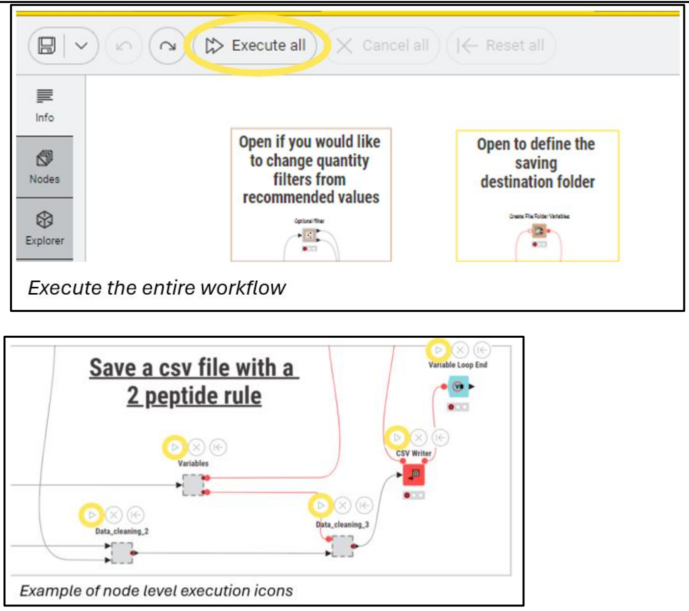

Figure DS2-6: Workflow execution

S2.7. <u>Visualization (Figure DS2-7)</u>

Open visual outputs at node level by selecting **Open View** (F10) on red component nodes. This launches interactive plots and tables within a composite visualization panel.

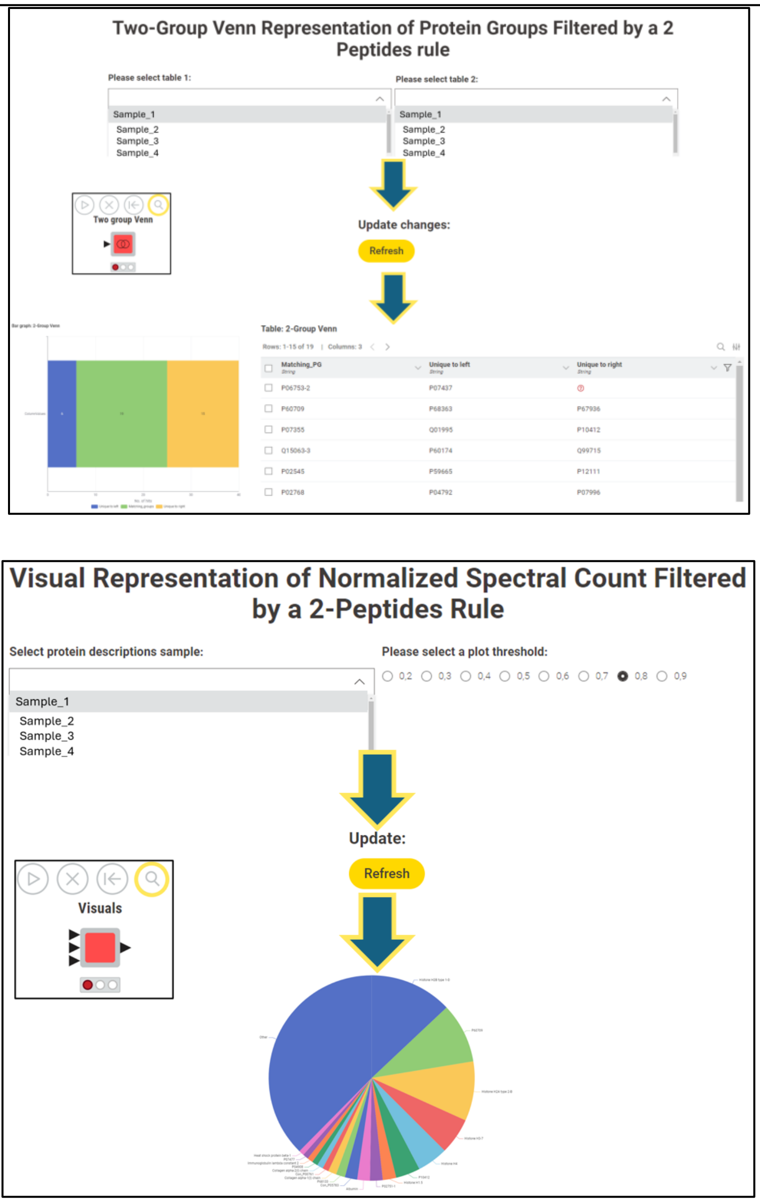

Figure DS2-7: Visualization

S2.8. <u>Grouped Visualizations (Figure DS2-8)</u>

Access "Grouped_Visuals" via **Open View**. Widget options allow adjustment of missing value thresholds and relative standard deviation (RSD) cut-offs per selected group. After refreshing, group-level summaries, intensity tables, and parallel coordinate plots are generated.

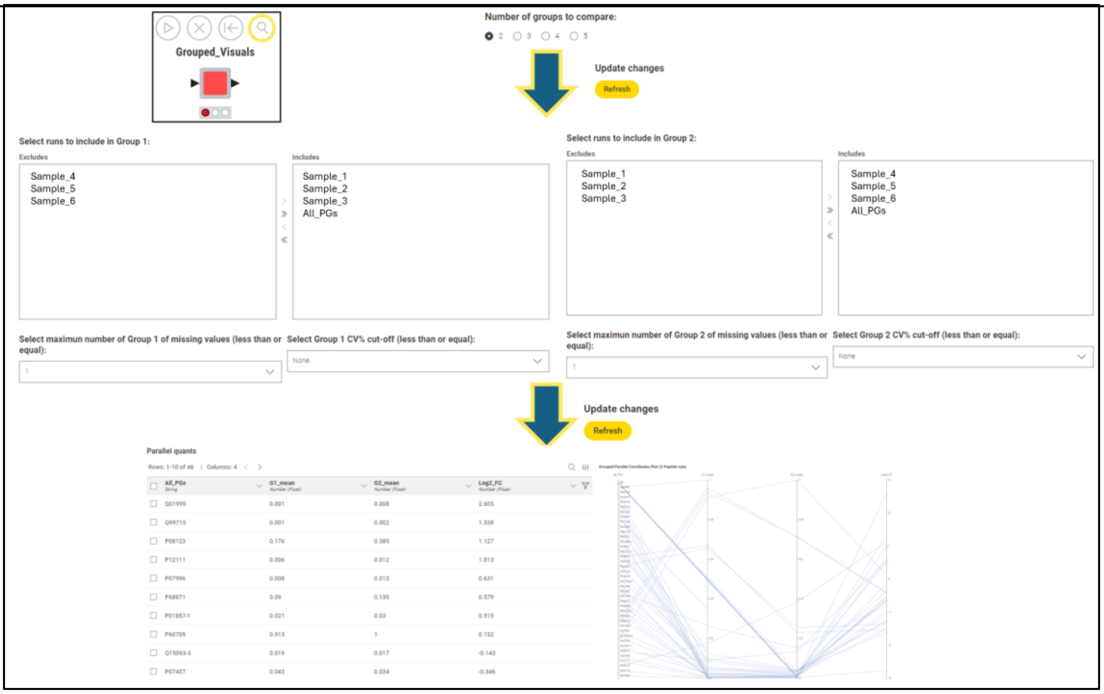

Figure DS2-8: Grouped Visualizations

S2.9. <u>Table Export (Figure DS2-9)</u>

Processed output tables are available at ports 1, 2, 5, 6, 7, and 8. Execute these nodes individually to export results for downstream bioinformatic or statistical analyses.

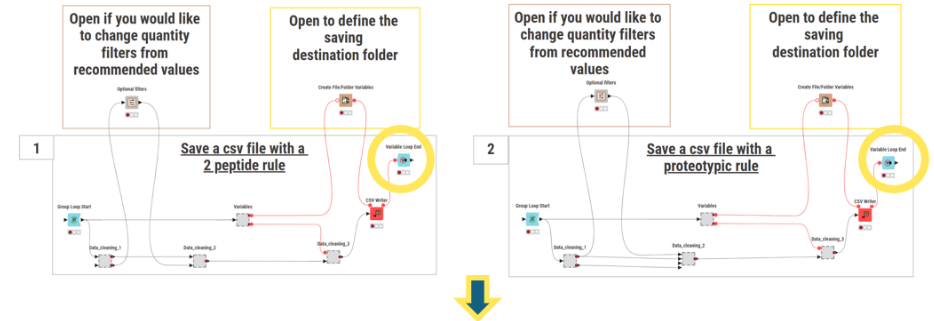

| First.Protein.Description | Sample_1_PG-ID | Unique_Stripped.Sequence | Sequence_count | Modifications | Modification_Count | Sample_1_Quant |
|---|---|---|---|---|---|---|
| ....... | ....... | ....... | ....... | ....... | ....... | ....... |
| ....... | ....... | ....... | ....... | ....... | ....... | ....... |

*Run-level csv output*

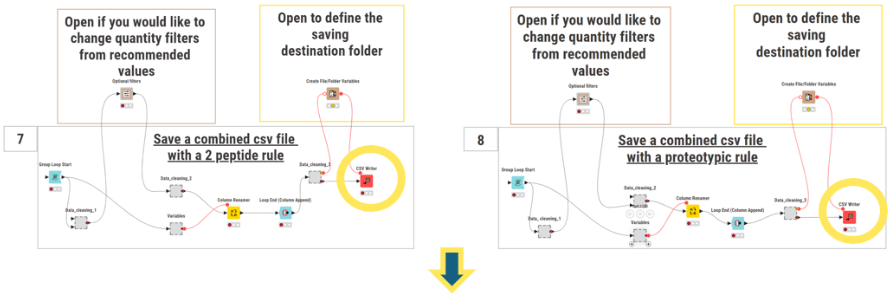

| All_PGs | Sample_1_Quant | Sample_2_Quant | Sample_3_Quant | Sample_4_Quant |
|---|---|---|---|---|
| ...... | ...... | ...... | ...... | ...... |
| ...... | ...... | ...... | ...... | ...... |

*Compound-level csv output*

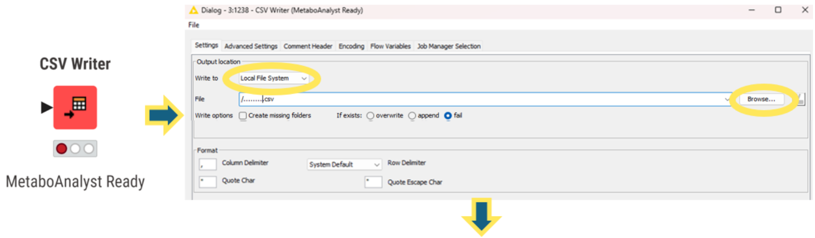

| All_PGs | Sample_1_Quant | Sample_2_Quant | Sample_3_Quant | Sample_4_Quant | Sample_5_Quant | Sample_6_Quant |
|---|---|---|---|---|---|---|
| Group | 1 | 1 | 1 | 2 | 2 | 2 |
| ..... | ..... | ..... | ..... | ..... | ..... | ..... |
| ..... | ..... | ..... | ..... | ..... | ..... | ..... |

*Group-level MetaboAnalyst-ready output*

Figure DS2-9: Table Export
